# Complete enzymatic depolymerization of polyethylene terephthalate (PET) plastic using a *Saccharomyces cerevisiae*-based whole-cell biocatalyst

**DOI:** 10.1101/2024.07.20.604433

**Authors:** Siddhant Gulati, Qing Sun

## Abstract

Plastics like polyethylene terephthalate (PET) have become an integral part of everyday life, yet plastic waste management remains a significant challenge. Enzymatic biocatalysis is an eco-friendly approach for recycling and upcycling of plastic waste. PET-hydrolyzing enzymes (PHEs) such as *Is*PETase, along with its engineered variants like FAST-PETase, demonstrate promising PET depolymerization capabilities at ambient temperatures. Whole-cell biocatalysts, displaying PHEs on their cell surface, offer high efficiency, reusability, and stability for PET depolymerization. However, their efficacy in fully breaking down PET is hindered by the necessity of two enzymes - PETase and MHETase. Current whole-cell systems either display only one PHE or struggle with performance when displaying larger passenger proteins like the MHETase-PETase chimera. In this work, we developed a *Saccharomyces cerevisiae*-based whole-cell biocatalyst system for complete PET depolymerization. Leveraging a cellulosome-inspired trifunctional protein scaffoldin displayed on the yeast surface, we immobilized FAST-PETase and MHETase, forming a multi-enzyme cluster. Our whole cell biocatalyst achieved complete PET depolymerization at 30°C, yielding 4.9 mM TPA in seven days with no intermediate accumulation. Furthermore, we showed improved PET depolymerization ability by binding FAST-PETase at multiple sites on the trifunctional scaffoldin. This breakthrough in complete PET depolymerization marks an essential step towards a circular plastic economy.

## Introduction

Polyethylene terephthalate (PET) plastic has become ubiquitous in daily life, finding extensive applications in the single-use packaging and textile industry. The annual global PET production is estimated to be around 82 million tonnes.^1^ However, since only a small fraction of this produced plastic is recycled (<20%), PET waste management is a daunting challenge.^2^ PET can be recycled via mechanical, chemical, or biological methods.^3,4^ Enzymatic biocatalysis is rapidly emerging as a green route for plastic waste recycling and could also open up new avenues for ‘upcycling’ waste into higher-value products.^5,6^.

Over the years, many PET-hydrolyzing enzymes (PHEs) have been identified and engineered for enzymatic PET recycling.^7,8^. The mesophilic enzyme, *Is*PETase, secreted by a gram-negative bacterium *Ideonella sakaiensis*, exhibited PET depolymerization ability at ambient temperatures.^9^ Engineered *Is*PETase variants with improved activity and thermostability have since been developed^10,11^, with FAST-PETase^12^ exhibiting superior performance than other engineered variants. Whole-cell biocatalysis is a possible approach to achieve PET depolymerization using enzymatic systems. By displaying PET-degrading enzymes on the surface of a prokaryotic or eukaryotic cell, promising PET depolymerization performance can be obtained.^13–18^ The benefits of these systems include improved activity due to reduced enzyme aggregation^14^, reusability, and better enzyme stability.

However, to achieve complete PET depolymerization into its constituent monomers-terephthalic acid (TPA) and ethylene glycol (EG), an enzymatic partner MHETase is also required.^9^ PETase and MHETase work synergistically to completely degrade PET, with MHETase-PETase fusion enzymes exhibiting even better performance than free enzyme mixtures.^19,20^ Additionally, the accumulation of intermediate product mono(2-hydroxyethyl) terephthalate (MHET) in the absence of MHETase has an inhibitory effect on enzymatic PET hydrolysis.^20–22^ A major drawback of existing PETase-based whole-cell biocatalyst systems is their ability to display only one enzyme efficiently on their surface.^13,14,16^ Poor performance^18^ or incomplete conversion^23^ is observed when larger passenger enzymes like the MHETase-PETase chimera are surface displayed. Thus, complete PET depolymerization remains challenging in such systems.

To achieve hydrolysis of recalcitrant polymers like cellulose in nature, certain anaerobic bacteria have developed multienzyme complexes called cellulosomes.^24^ Inspired by these natural systems, artificial, designer cellulosomes have been developed to harness the benefits of enzyme proximity, substrate channeling, and synergy, which lead to highly efficient substrate degradation.^25,26^

In this work, we developed a *Saccharomyces cerevisiae*-based whole-cell biocatalyst system capable of complete PET depolymerization. Previous research has demonstrated the excellent surface display capability of *S. cerevisiae*-based systems.^27,28^ Additionally, the scaffold approach provides flexibility, eliminating the need for exhaustive experimental screening to surface display enzymes, thus rendering it highly direct.^29–31^ FAST-PETase and MHETase were bound on a trifunctional protein scaffoldin displayed on the yeast cell surface through high-affinity dockerin-cohesin interactions. We confirmed the scaffoldin’s display on the yeast cell’s surface and the successful binding of each enzyme onto this scaffoldin. Subsequently, we assessed the PET depolymerization activity of this whole-cell biocatalyst and showcased its capability to completely break down PET into its constituent monomers, TPA and EG. This work is essential to achieve a robust and eco-friendly plastic waste management approach.

## Materials and Methods

### Yeast culture and display of trifunctional scaffoldin on the yeast cell surface

*S. cerevisiae* strain EBY100 [*MAT*a *AGA1*::*GAL1-AGA1*::*URA3 ura3*-*52 trp1 le*u*2*-*1 his3*-*200 pep4*::*HIS3 prb1*-*1*.*6R can1 GAL*] transformed with pScaf-ctf^32^ was used for the surface display of the trifunctional scaffoldin.

The yeast cells harboring the surface display plasmid were precultured in SDC medium (20g/L dextrose, 5g/L Bacto casamino acids, 6.7g/L yeast nitrogen base without amino acids) at 30°C for 18h. For the surface display of the trifunctional scaffoldin, the precultured cells were inoculated into SGC medium (20g/L galactose, 5g/L Bacto casamino acids, 6.7g/L yeast nitrogen base without amino acids) at OD_600_ of 0.5. These were subsequently allowed to grow at 20°C for 60h. *S. cerevisiae* EBY100 without the pScaf-ctf plasmid served as a negative control.

### Expression and purification of PET-degrading enzymes

Sequences of gene fragments encoding for dockerins from *C. cellulolyticum* (Doc Cc), *C. thermocellum* (Doc Ct), and *R. flavefaciens* (Doc Rf) were obtained from a previous publication.^32^ Dockerin-tagged FAST-PETase and MHETase were constructed by fusing the dockerins at the C-terminus of the protein of interest (POI) separated by a flexible GS linker and transformed into *E. coli* NEB 5-alpha.

The constructed plasmids, pBbE8k-FAST-PETase-Dockerin Rf (pBB8-FP-Rf), pBbE8k-FAST-PETase-Dockerin Cc (pBB8-FP-Cc) and pBbE8k-MHETase-Dockerin Ct (pBB8-MH-Ct) were subsequently transformed into *E. coli* SHuffle T7 for protein expression. The cells harboring pBB8-FP-Rf and pBB8-FP-Cc were inoculated into fresh LB medium containing kanamycin, cultured overnight at 37°C, and then scaled up at the same temperature until OD_600_ reached 0.9. Subsequently, intracellular protein expression was induced with 10 mM L-arabinose at 20°C/250RPM for 20h. The cells harboring pBB8-MH-Ct were grown in TB medium and protein expression was induced with 1 mM L-arabinose.

Post-protein expression, the cells were lysed and the proteins were purified by His tag purification (HisPur Ni-NTA Resin, Thermo Scientific), followed by concentration and buffer exchange into Exchange buffer (50mM HEPES, 100mM NaCl, pH 8) using a centrifugal filter. All purification steps were performed at 4°C. Protein purity was assessed using SDS-PAGE and quantified using Bio-Rad Image Lab software. Protein concentration was determined using Bio-Rad DC Protein Assay.

### Enzyme assembly on the yeast cell surface

Yeast cells displaying the scaffoldin were resuspended in Binding buffer (50 mM Tris HCl, 100 mM NaCl, 10 mM CaCl_2_, pH 8). Subsequently, purified dockerin-tagged enzymes were added and incubated with the yeast cells at 20°C for 1.5h with continuous shaking. After incubation, the yeast cells were washed with 1x PBS to remove unbound dockerins, and resuspended in Binding buffer for activity assays.

### Enzyme activity assays on PET

Amorphous PET film (Goodfellow, ES301445) was used as substrate to test the PET hydrolytic activity of the whole-cell biocatalyst. PET films were hole-punched into circular films (6 mm diameter) and thoroughly washed with 1% SDS, 20% ethanol, and deionized water prior to use. The PET films were added to a glass test tube containing 300μL of yeast cells (OD_600_ 10) followed by incubation at 30°C and 250 RPM. Samples were collected periodically to quantify the reaction products.

To terminate the enzyme reaction, the reaction mixture was centrifuged at 10000g for 5 minutes to remove yeast cells, followed by heat treatment of the reaction supernatant at 85°C for 20 minutes. This reaction supernatant was subsequently analyzed using HPLC (detection wavelength 240 nm) to quantify the formation of MHET and TPA by comparing against a standard curve prepared using commercially available standards.

### Confirmation of scaffoldin surface display and determination of FAST-PETase saturation concentration

To confirm the surface display of the scaffoldin, *S. cerevisiae* cells displaying the scaffoldin (OD_600_ 1) were harvested by centrifugation and washed with 1x PBS. The cells were resuspended in 100μL Blocking buffer (1xPBS containing 1 mg/ml bovine serum albumin) and left on a rotary nutator for 1h for blocking. This was followed by incubation with the primary anti-C-myc antibody (Myc-Tag (71D10) Rabbit mAb #2278, Cell Signaling Technology, 1:200 dilution) at 4°C overnight. The cells were then washed with 1x PBS and resuspended in blocking buffer, followed by the addition of the secondary antibody conjugated to Alexa 488 (Anti-rabbit IgG (H+L), F(ab’)2 Fragment (Alexa Fluor® 488 Conjugate) #4412, Cell Signaling Technology, 1:500 dilution) at room temperature for 2h in the dark on a rotary nutator. After incubation, the cells were washed with 1x PBS and fluorescence was measured using a microplate reader (BioTek Synergy H1) at the following wavelengths: λ_ex_ = 488 nm, λ_em_ = 530nm.

To determine saturation concentration of FP-Rf and FP-Cc on the scaffoldin, equimolar amounts of both enzymes were bound to the scaffoldin using the enzyme assembly protocol. *S. cerevisiae* cells (OD_600_ 5) containing the bound dockerins were centrifuged and washed, followed by blocking for 1h in Blocking buffer. They were then incubated with the anti-C-His antibody conjugated to Alexa Fluor 488 (His-Tag (D3I1O) XP® Rabbit mAb (Alexa Fluor® 488 Conjugate) #14930, Cell Signaling Technology, 1:100 dilution) for 1h at room temperature in the dark on a rotary nutator. After incubation, the cells were washed with 1x PBS and fluorescence was measured (λ_ex_ = 488 nm, λ_em_ = 530nm).

## Results and discussion

### Functional display of scaffoldin on the surface of the yeast cell

A trifunctional scaffoldin, comprised of three cohesin domains from *C. thermocellum* (Coh Ct),

*C. cellulolyticum* (Coh Cc), and *R. flavefaciens* (Coh Rf) with a cellulose-binding domain (CBD) was displayed on the surface of the *S. cerevisiae* cell using the Aga1-Aga2 anchor system (Figure 1A).^32^ To verify the surface display of the scaffoldin, a c-myc tag on the scaffoldin was utilized. Upon incubating *S. cerevisiae* whole-cells with fluorophore-conjugated anti-c-myc antibody, considerably higher fluorescence intensity was observed in cells displaying the scaffoldin compared to wild-type whole-cells (Figure 1B), confirming the surface localization of the scaffoldin.

**Figure 1.**
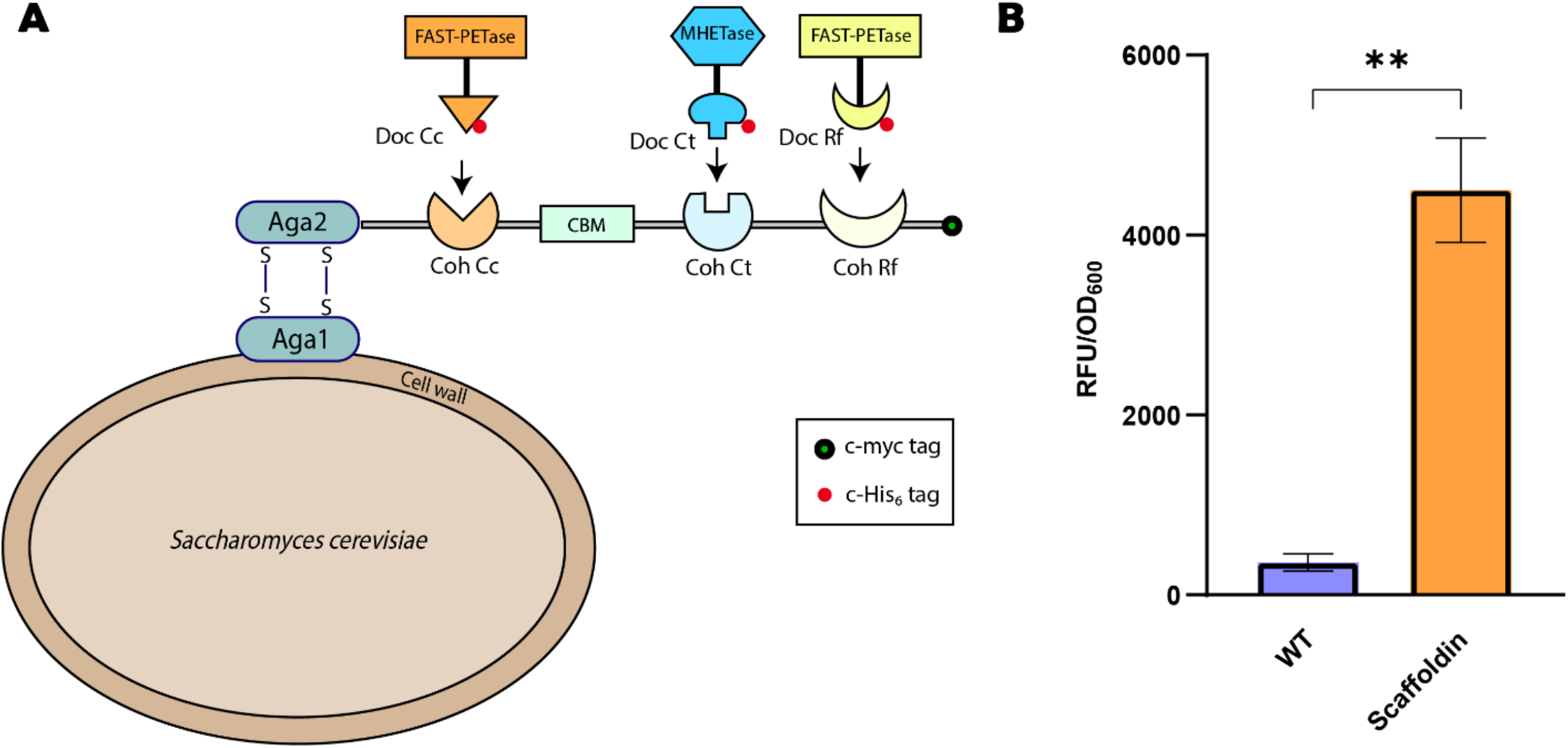
**(A)** Schematic representation of trifunctional scaffoldin displayed on the yeast cell surface. The scaffoldin consists of cohesin domains derived from *C. cellulolyticum* (Cc), *C. thermocellum* (Ct) and *R. flavefaciens* (Rf) and an internal cellulose-binding domain (CBD). Dockerin-tagged FAST-PETase (FP) and MHETase (MH) bind specifically to their corresponding cohesin domains; **(B)** Fluorescence intensity of scaffoldin-displaying whole cells probed using a primary anti-C-myc antibody and an Alexa Fluor 488-conjugated secondary antibody. Wild-type (WT) yeast cells served as negative control. Experiments were conducted in triplicates, data shown are mean values (± standard deviation). Statistical significance was evaluated using unpaired student *t*-tests. \*\**P*<0.01.

### Binding of PET-degrading enzymes on the scaffoldin

Previous studies have shown that PETase and MHETase work synergistically to completely degrade PET. It has been reported that higher PETase loading leads to high degradation performance and even low MHETase concentrations are sufficient to boost product release and completely convert MHET to TPA.^19^ Based on this understanding, we hypothesized that attaching FAST-PETase at two different sites would allow higher FAST-PETase concentrations than single-site attachment of FAST-PETase, promoting efficient and complete PET depolymerization. In our approach, we utilized FAST PETase-Doc Rf (FP-Rf) and FAST PETase-Doc Cc (FP-Cc) for immobilizing FAST-PETase on the surface scaffoldin and MHETase-Doc Ct (MH-Ct) for MHETase immobilization (Figure 1).

We determined the saturation concentration of FAST-PETase on the scaffoldin by incubating equimolar concentrations of FP-Rf and FP-Cc with the yeast cells (Figure S1). This concentration of the FAST-PETase dockerin enzymes was used for activity assays on the PET film in this work.

Meanwhile, when wild-type yeast cells (lacking the scaffoldin) were incubated with all three enzymes, we observed negligible TPA formation compared with scaffoldin display cells. This observation confirmed minimal non-specific enzyme binding on the yeast surface in the absence of the scaffoldin (Figure 2).

**Figure 2.**
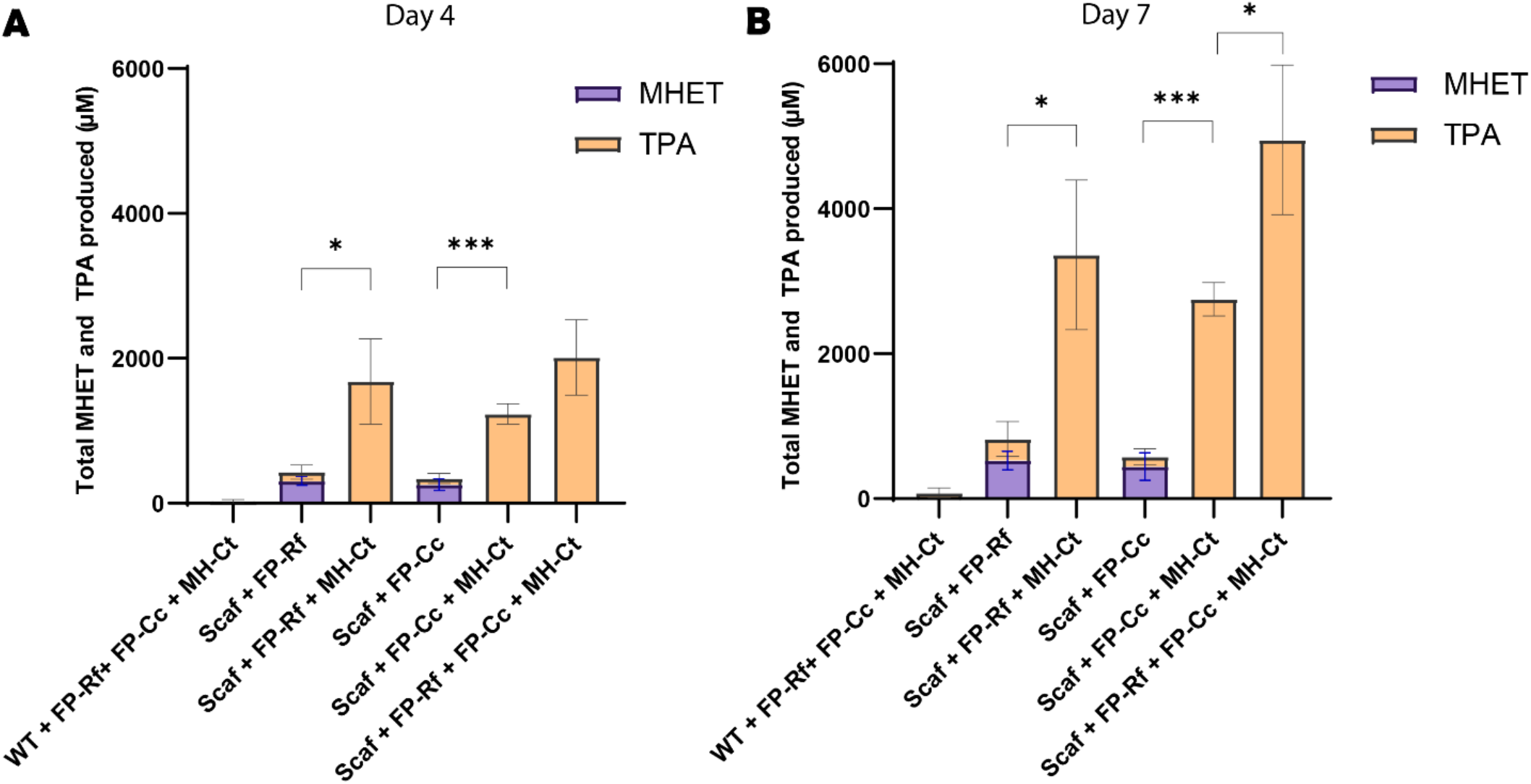
Determination of PET hydrolytic activity of whole-cell biocatalyst, evaluated by quantification of amount of MHET and TPA released in reaction supernatant via HPLC after degradation of PET film at **(A)** Day 4 and **(B)** Day 7. Reactions were performed at 30°C in Binding buffer. (Whole-cell OD_600_=10, FP-Rf = 400 nM, FP-Cc = 400 nM, MH-Ct = 50 nM). Experiments were conducted in triplicates, data shown are mean values (± standard deviation). Statistical significance was evaluated using unpaired student *t*-tests. \**P*<0.05, \*\*\**P*<0.001.

### Study of PET depolymerization performance of whole-cell biocatalyst

Following incubation of *S. cerevisiae* whole cells with the PET film, we quantified reaction products in the supernatant using HPLC to monitor MHET and TPA formation. When only FAST-PETase dockerins were present on the scaffoldin, both MHET and TPA were formed (Figure 2, Figure S2).

When MH-Ct was present on the scaffoldin along with FP-Cc and FP-Rf, we observed complete PET depolymerization, yielding only TPA as the enzymatic reaction product, with no MHET accumulation detected (Figure 2, Figure S2). This demonstrates that each enzyme, including PETase and MHETase, was active after binding to the scaffoldin, and the PETase-MHETase pair was able to completely depolymerize PET with no intermediate accumulation. As expected, MH-Ct alone did not show any activity on the PET film (data not shown).

On Day 4, the scaffoldin-displaying yeast cells bound with FP-Rf and MH-Ct exhibited better performance than their FP-Cc and MH-Ct counterpart (Figure 2A). This difference in performance could be attributed to variance in enzyme localization and access to plastics. We also explored the effect of having FAST-PETase at two sites (with MHETase at one site) on the scaffoldin, aiming to further enhance the concentration of FAST-PETase in the depolymerization reaction. The three-enzyme system exhibited up to 1.63 times better depolymerization performance than the two-enzyme counterparts throughout Day 4, with mean TPA yields of 2013 µM.

It was observed that the total product formation was significantly boosted when MH-Ct was co-immobilized with FP-Rf and FP-Cc on the scaffoldin (Figure 2A, 2B). This could be attributed to the inhibitory effect of MHET accumulation in the absence of MHETase, which has been shown to have a detrimental impact on PETase activity.^20,21^

Throughout Day 7 (Figure 2B), the whole cells remained active, with at least a 2-fold increase in TPA production relative to Day 4. Notably, binding of FAST-PETase at multiple sites (with MHETase at one site) on the scaffoldin proved extremely beneficial, resulting in a 1.4-fold increase relative to FP-Rf (with MH-Ct) and a 1.8-fold increase relative to FP-Cc (with MH-Ct). A mean yield of 4.9 mM TPA was obtained after 7 days of incubation with the PET film when all three enzymes were bound to the scaffoldin. With these results, we successfully demonstrated the advantages of immobilizing FAST-PETase at multiple sites on the scaffoldin for efficient and complete PET depolymerization.

## Conclusion

We developed an *S. cerevisiae* based whole-cell biocatalyst capable of fully depolymerizing PET into its constituent monomers, TPA and EG. The PET-degrading enzymes, FAST-PETase and MHETase, were immobilized on the surface of the yeast cell by binding to a trifunctional scaffoldin displayed on the yeast surface in a highly site-specific manner. TPA yields of 4.9 mM were achieved in 7 days at 30°C, with no observed accumulation of reaction intermediates. The attachment of FAST-PETase at two sites on the trifunctional scaffoldin proved advantageous, as PETase is required in excess compared to MHETase in the enzymatic PET depolymerization process.

The produced TPA and EG can be reused by both chemical and biological methods. They can facilitate the regeneration of virgin PET, enabling closed-loop PET recycling.^12^ Additionally, these products can be repurposed into various value-added chemicals and bioplastics using alternative biocatalysts.^2,33,34^ Thus, this PET depolymerization approach could pave the way for a circular plastic economy.

## Declaration of Competing Interest

The authors declare that they have no known competing financial interests or personal relationships that could have appeared to influence the work reported in this paper.

### Credit Author Statement

Q. S. did the conceptualization. S. G. designed the experiment, conducted experiment, and drafted the manuscript. Q. S. revised the manuscript and did the funding acquisition.

## Acknowledgements

This work was supported by funds provided by a National Science Foundation (NSF) grant to Q. S. (2203715), USA; NSF Emerging Frontiers in Research and Innovation (EFRI) to Q. S. (2132156), USA; Texas A&M Excellence Fund X-grants to Q. S., Texas A&M, USA; Texas A&M Targeted Proposal Teams (TPT) grant to Q. S., Texas A&M, USA. The authors thank Dr. Wilfred Chen for providing the yeast strains.

## Supplementary information

**Figure S1:**
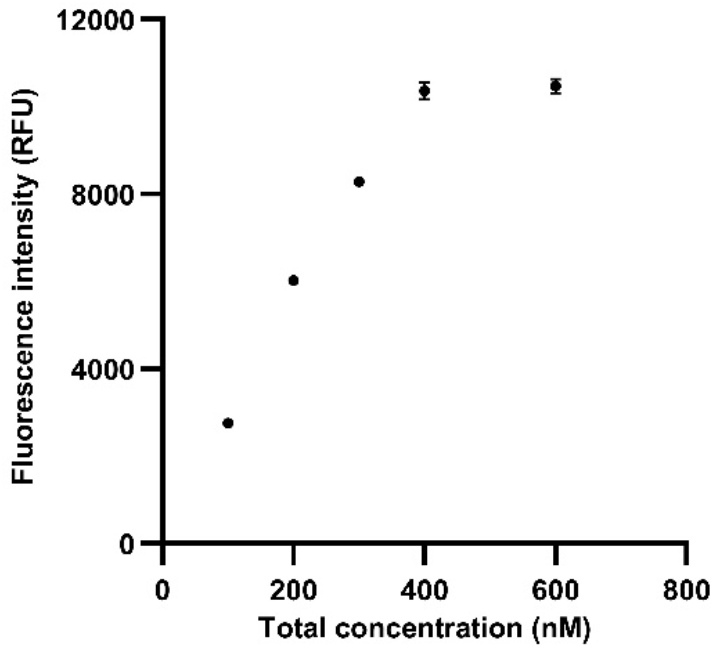
Determination of saturation concentration of yeast scaffoldin (OD_600_ 5) after incubation with equimolar concentrations of FP-Cc and FP-Rf. The x-axis represents total FAST-PETase concentration. Fluorescence intensity was determined after incubation with anti-C-His antibody conjugated to Alexa Fluor 488. Experiments were conducted in triplicates, data shown are mean values (± standard deviation). Error bars smaller than size of symbol are not visible.

**Figure S2:**
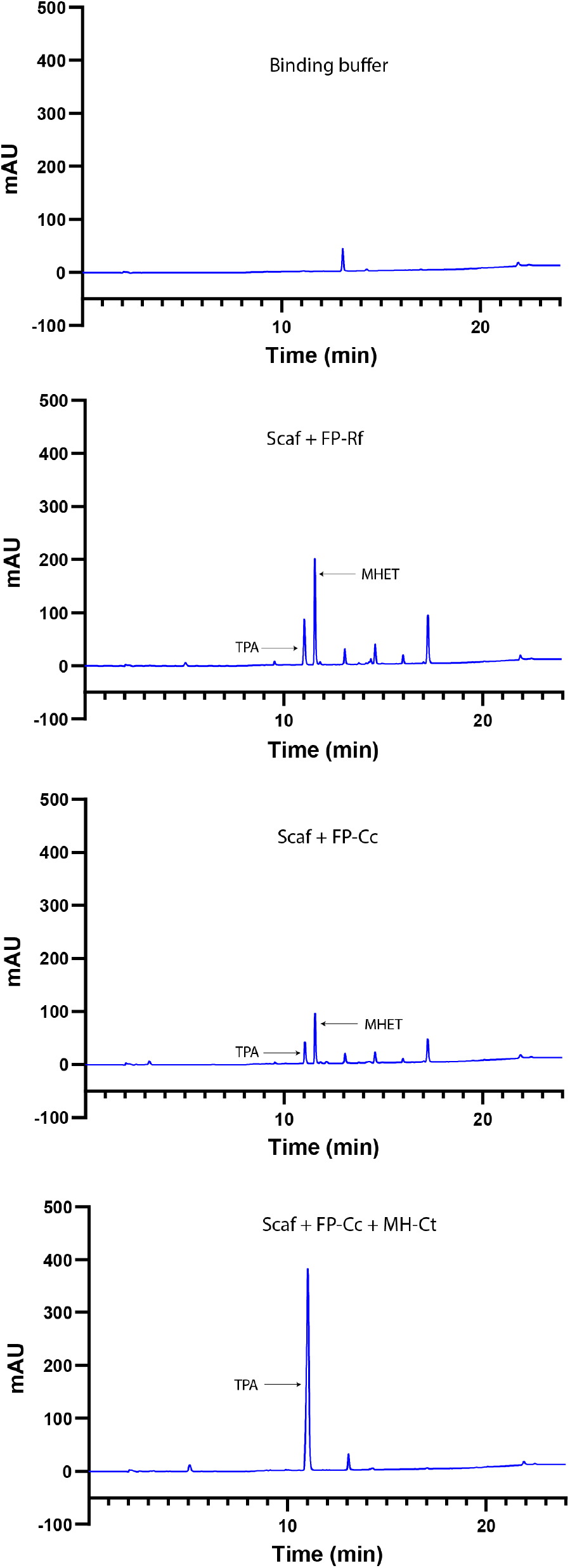
HPLC chromatogram of PET degradation products at Day 4. All experiments were conducted using Binding buffer as reaction medium.

